# Interactions of a bacterial RND transporter with a transmembrane small protein in a lipid environment

**DOI:** 10.1101/813394

**Authors:** Dijun Du, Arthur Neuberger, Mona Wu Orr, Catherine E. Newman, Pin-Chia Hsu, Firdaus Samsudin, Andrzej Szewczak-Harris, Leana M. Ramos, Mekdes Debela, Syma Khalid, Gisela Storz, Ben F. Luisi

## Abstract

The small protein AcrZ in *Escherichia coli* interacts with the transmembrane portion of the multidrug efflux pump AcrB and increases the resistance of the bacterium to a subset of the antibiotic substrates of that transporter. It is not clear how the physical association of the two proteins selectively changes activity of the pump for defined substrates. Here, we report cryo-EM structures of AcrB and the AcrBZ complex in lipid environments, and comparisons suggest that conformational changes occur in the drug binding pocket as a result of AcrZ binding. Simulations indicate that cardiolipin preferentially interacts with the AcrBZ complex, due to increased contact surface, and we observe that the drug sensitivity of bacteria lacking AcrZ is exacerbated when combined with cardiolipin deficiency. Taken together, the data suggest that AcrZ and lipid cooperate to allosterically modulate the activity of AcrB. This mode of regulation by a small protein and lipid may occur for other membrane proteins.

## Introduction

Numerous small proteins, corresponding to 100 or fewer codons, are encoded by phylogenetically diverse organisms and are likely to play key roles in many fundamental biological processes (Storz *et al*, 2014). Some small proteins have been discovered to regulate activity of large transport proteins, and in *Escherichia coli*, the 49 amino acid AcrZ modulates the action of AcrB, a homotrimeric secondary-active transporter that belongs to the resistance-nodulation-cell division (RND) superfamily (Hobbs *et al*, 2012). Deletion of the *acrZ* gene renders *E. coli* cells more sensitive to a subset of the antibiotics for which AcrB provides resistance (Hobbs *et al*, 2012). Transcription of *acrZ* is co-regulated with the *acrAB* operon, which also implicates the functional importance of the small protein for efflux activity (Hobbs *et al*, 2012).

AcrB is the energy-transducing component of a tripartite multi-drug efflux machinery that includes the outer membrane protein TolC and the periplasmic bridging partner AcrA. Structures of the fully assembled tripartite complex together with AcrZ have been elucidated using cryo-electron microscopy (Du *et al*, 2014; Jeong *et al*, 2016; Wang *et al*, 2017b), revealing that the small protein forms a transmembrane helix which interacts extensively with the concave surface of AcrB in the transmembrane region. The interaction of AcrZ and AcrB *in situ* has been corroborated by mass spectrometry of the intact complex ejected directly from native membranes of *E. coli* cells (Chorev *et al*, 2018).

While its influence over AcrB remains unclear from the available data, AcrZ was hypothesized to alter the conformation of the drug binding pockets during the transport cycle and so change drug specificity. In this model, AcrZ could exert an influence on AcrB by changing the surface exposed to the lipid from a concave to a convex curvature, thus potentially affecting the interactions with lipids and distribution of lateral forces of the bilayer that can be communicated into the core of the transporter. In support for this proposal, experimental findings in other systems indicate that lipids and the membrane composition can have profound effects on structure, oligomerization and activity of membrane proteins (Bechara *et al*, 2015; Gupta *et al*, 2017; Laganowsky *et al*, 2014). Moreover, a recent cryo-EM study of AcrB extracted directly from membranes reveals a semi-crystalline lipid organization within the central region of the transmembrane domains of the AcrB trimer that may support quaternary state transitions required for the transport mechanism (Qiu *et al*, 2018).

To investigate how AcrZ impacts AcrB, we determined cryo-EM structures of AcrB and the AcrBZ complex reconstituted in a disc in which a bilayer of *E. coli* lipids is encircled by the membrane scaffold protein saposin A (Denisov & Sligar, 2016). To facilitate particle alignments for 3D reconstructions, the complexes included an engineered DARPin protein that binds the periplasmic domain of AcrB (Eicher *et al*, 2012; Sennhauser *et al*, 2006). Trimeric AcrB cycles through three states in the transport process, and DARPin associates with the subunits in the loose (L) and tight (T) states, but not with the periplasmic region of the open state (O). These three states can be observed in the cryo-EM reconstructions, enabling analysis of the AcrZ interactions with each state. We conclude that the combination of AcrZ and lipid environment work synergistically to provide an allosteric effect on the conformation of AcrB, with functional consequences for the dynamic substrate transport process.

## Results

### Cryo-electron microscopy of DARPin-bound AcrB and AcrBZ reconstituted into saposin A discs

We developed a procedure to reconstitute purified *E. coli* AcrB and the AcrBZ complex into discs using *E. coli* lipids and saposin A as scaffolding protein (details in Materials and Methods). The reconstituted specimens eluted with a gausian shaped profile behaved well on size exclusion chromatography in buffer without detergent that is otherwise required to keep the membrane proteins soluble. To facilitate particle alignment from cryo-EM images of these specimens, we included an engineered DARPin that binds the periplasmic domain of AcrB (Eicher *et al*, 2012; Sennhauser *et al*, 2006). The disc-reconstituted, DARPin-bound AcrB and AcrBZ samples yielded excellent quality particles on cryo-EM grids (Appendix Fig S1A and S1B). Analysis of the particles provided interpretable maps with resolutions near 3.2 Å based on Fourier shell correlations. Models could be built into the density and refined with good stereochemistry (Appendix Table S1). Top, bottom and side views of AcrB and AcrBZ are shown in Fig 1.

**Figure 1.**
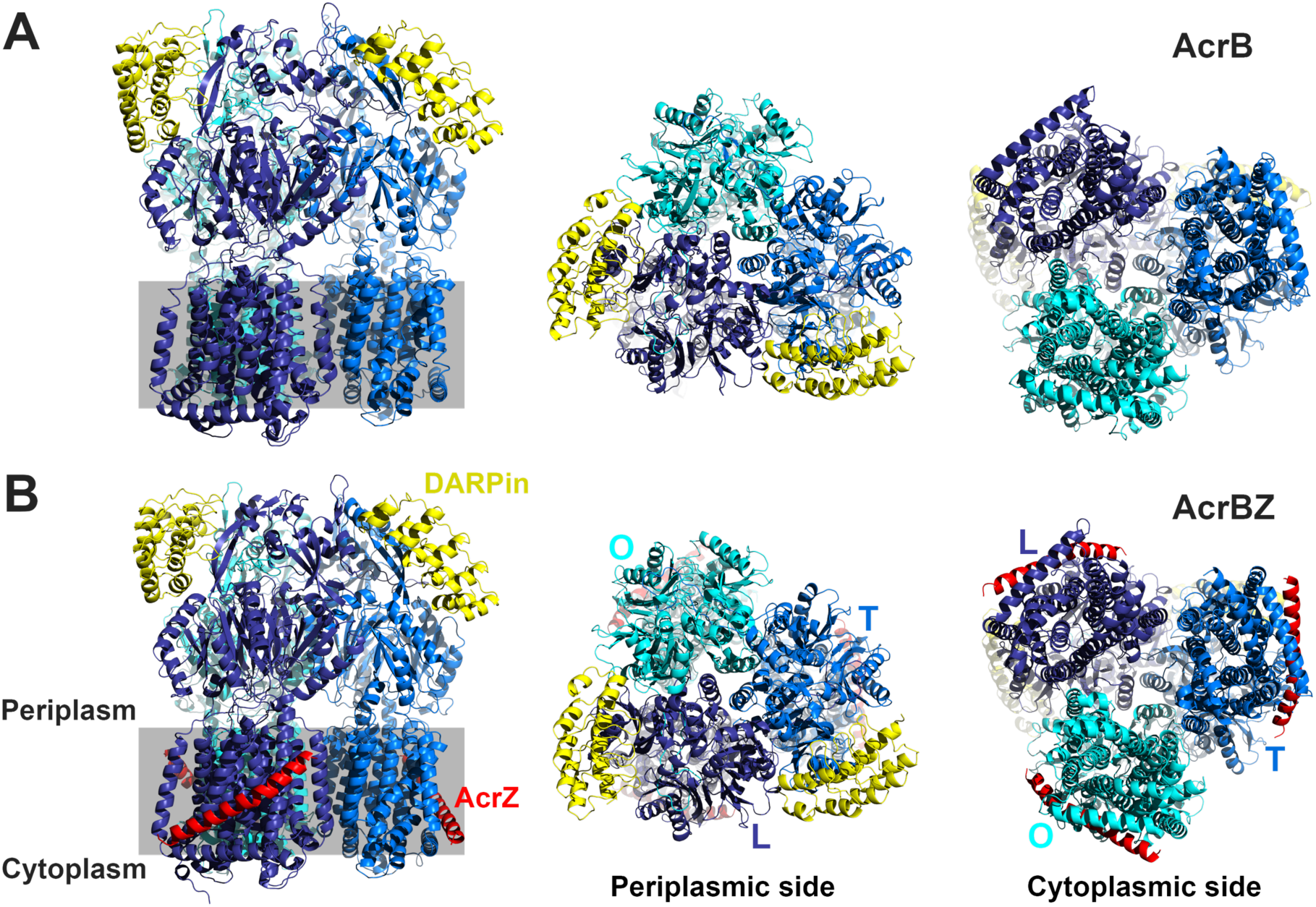
*E. coli* AcrB and AcrBZ in saposin A discs. A Structure of AcrB in saposin A discs prepared using *E. coli* lipids: Side (left; oriented so that the three-fold axis is aligned vertically), periplasmic (center; along the three-fold axis), and cytoplasmic (right; along the three-fold axis) view. Within the trimer, the AcrB subunits are found in loose (L, navy blue), tight (T, blue) and open (O, cyan blue) conformations. Two DARPin domains (yellow) are positioned at the top of the periplasmic domain; attached to protomers in loose (L) and tight (T) conformations. B Refined model of AcrBZ fitted to the cryo-EM map in three views (see above). Two DARPin domains (yellow) are attached to protomers in L and T. AcrZ is displayed in red.

Saposin A monomers, which form a ring around the transmembrane domain of AcrB and the AcrBZ complex, were also resolved at a lower density, indicating that the ring is associated flexibly with the transporter. The features of the saposin A become more diffuse as refinement progressed, while the features of the AcrB and AcrBZ become sharper. A lipid layer could be visualized in the maps, and acyl chains or lipid head groups were included in the refined models.

A well-organised lipid bilayer within the central region of the transmembrane domains of the AcrB trimer was noted in earlier studies of specimens solubilised with styrene-maleic acid co-polymers (Qiu *et al*, 2018). We also observe a lipid layer in our maps in the corresponding region (Fig 2A). Our best model for the AcrBZ complex in saposin A discs is consistent with acyl chain packing observed in the structure of AcrB in co-polymers (Fig 2A). Simulations show that the acyl chains on the inner leaflet pack in a stable lattice with approximate hexagonal geometry (Fig 2B), visible also in our cryo-EM maps, while they form a less regular pattern in the outer leaflet. The calculations indicate that acyl chains are more mobile in the outer layer compared to the inner layer (Fig 2C). Hydrophobic side chains of AcrB are predicted to interact with the acyl groups of these lipids in different ways for the three protomers (Fig 2D); such differences could help to communicate conformational signals between the subunits associated with transitions between the L, T, and O states, as proposed earlier (Qiu *et al*, 2018). Density is also present for lipids on the outer surface where AcrZ interacts, but the lipids here are not extensively ordered.

**Figure 2.**
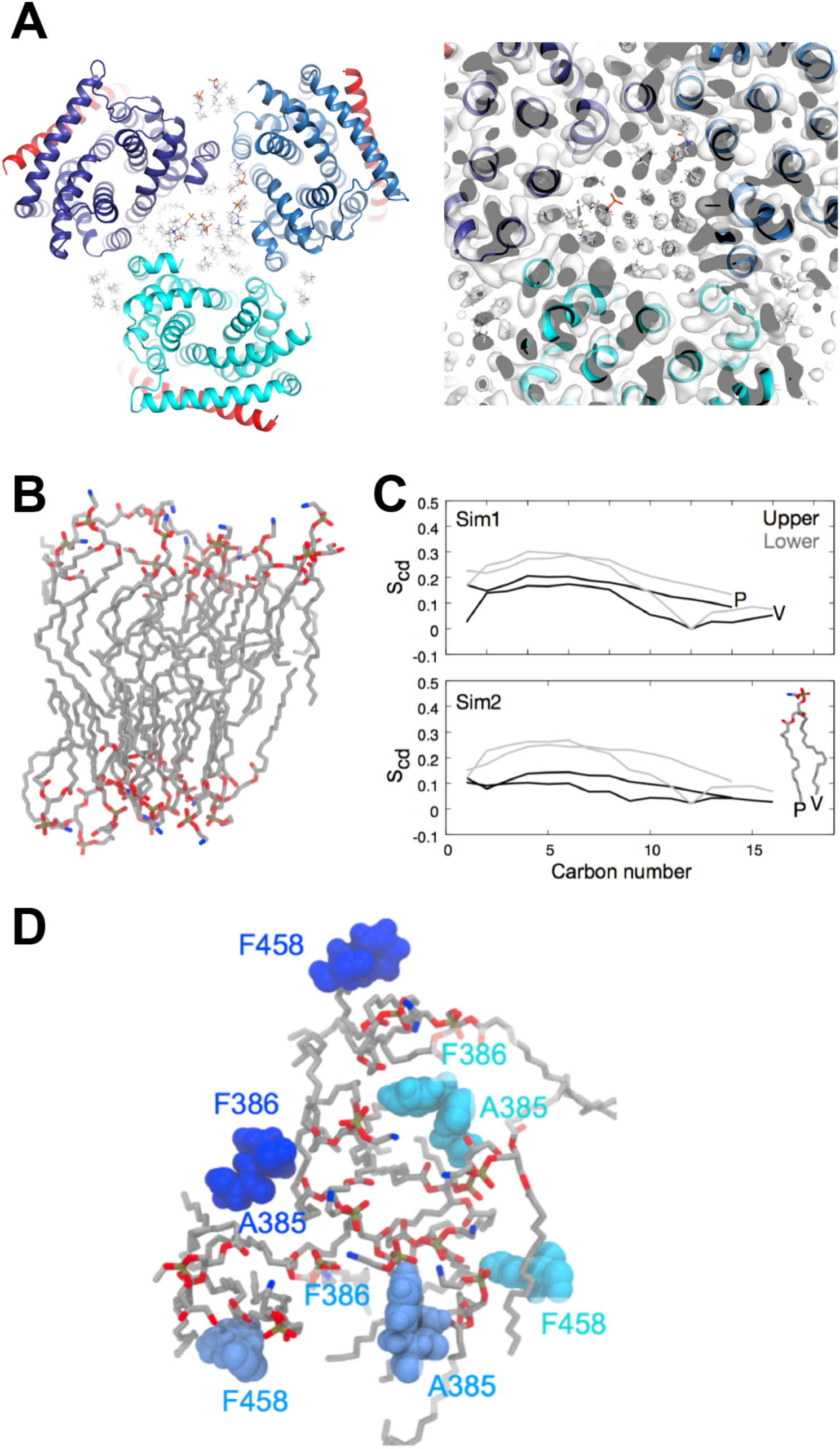
Organization and interactions of lipids in the central cavity of AcrB. A Left panel shows a view along the central axis of the AcrBZ complex with *E. coli* lipids at natural abundance. Right panel shows the cryo-EM density in the central region which reveals the hexagonal pattern for the lipids. Colours of AcrB subunits and AcrZ are the same as in Fig 1. B Snapshots of the lipid bilayer in the central cavity of AcrB at the end of a 500 ns simulation. C Average order parameters (S_cd_) of the palmitic (P) and vaccenic (V) acid lipid tails of PVPE in the outer and inner leaflets of the central cavity in two simulations (Sim1 and Sim2). D Hydrophobic residues that protrude into the outer leaflet of the lipid bilayer during the simulations in the different AcrB protomers (L, navy blue; T, blue; and O, cyan blue).

### The lipid environment affects the orientation of AcrZ on AcrB

The models of AcrB and AcrBZ in the saposin discs were compared against previously described crystal structures of detergent-solubilised AcrB and AcrBZ, respectively (Fig 3). As a reference frame for the comparison, we used transmembrane helices (TMH) 4-6, which had previously been demonstrated to be a suitable group for overlays due to its conformationally invariance in the L, T and O states (Murakami *et al*, 2006; Seeger *et al*, 2006). Small conformational shifts were detected between the detergent-based crystal structure and the saposin disc-reconstituted cryo-EM structure for both AcrB and AcrBZ, including a rotation of the periplasmic domain (foremost in PC2, PN2 and PC1 sub-domains) in all three states (L, T and O). These differences in the periplasmic domain could reflect the response of the protein to constraints imposed or enforced by the crystal lattice. A structural comparison of AcrB and AcrBZ when both are in saposin A discs indicates small changes in all three AcrB protomers in PC1/2 and PN2 and TMH2 and 8 (Appendix Fig S2). These changes may be a consequence of the presence of AcrZ.

**Figure 3.**
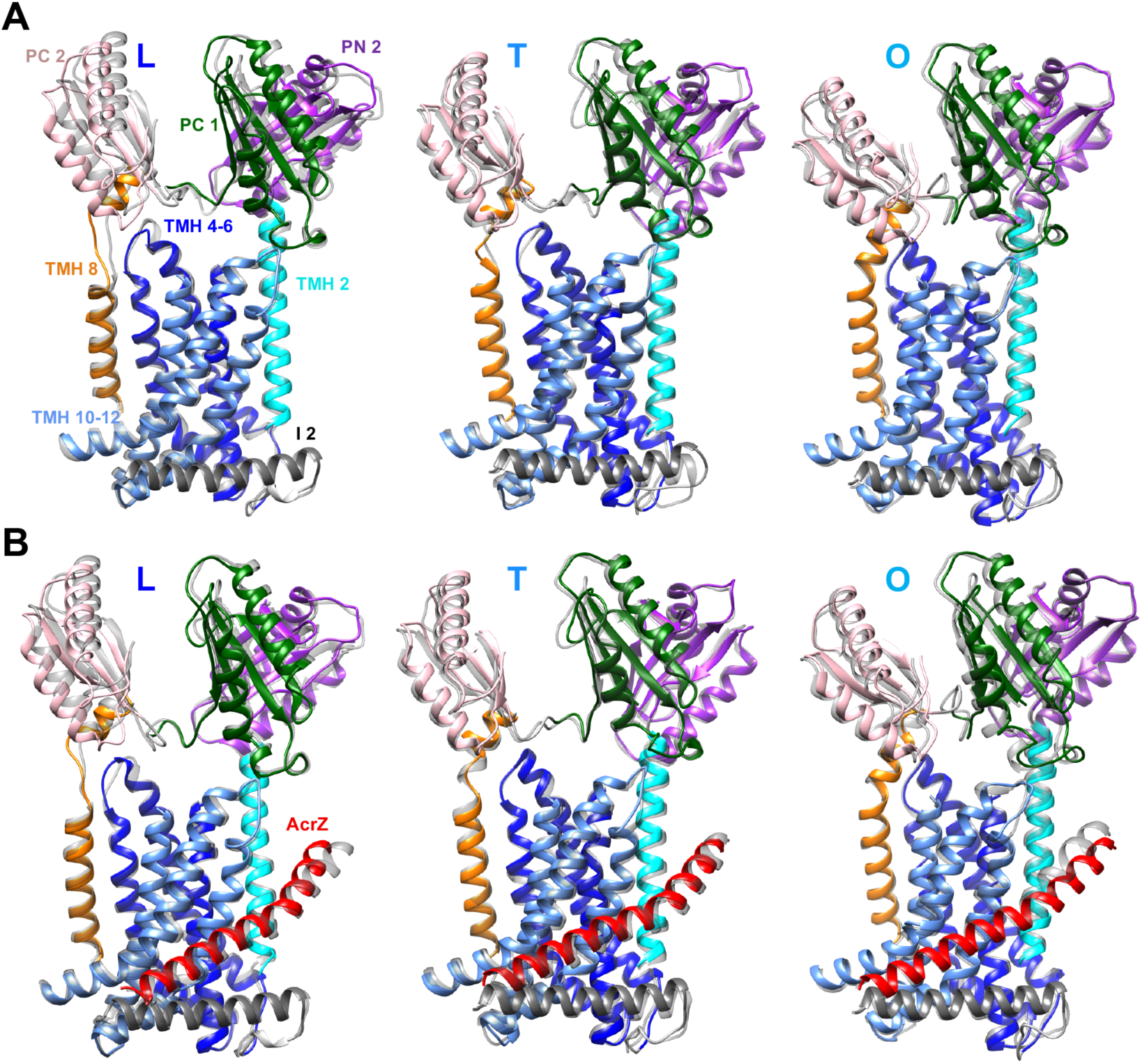
Gallery of structural comparisons of AcrB and AcrBZ crystal structures (grey) overlaid by cryo-EM saposin A-nanodisc structures (coloured). A Overlay of protomers in L, T, O of (1) AcrB crystal structure (PDB-ID: 4DX5) (light grey; partially transparent) and (2) cryo-EM derived AcrB structure reconstituted in *E. coli* lipids inside a saposin A-nanodisc. Orientations of protomer overlays L, T, O in space were adjusted in depiction for a better display of observed changes. Colour Code: PC2, pink; PN2, purple; PC1, dark green; TMH 8, orange; TMH 4-6, navy blue; TMH 2, cyan blue; TMH 10-12, blue; I2, grey. Displayed are PC1/2, PN2, I2, TMH 2, TMH 8, TMH 10-12, and TMH 4-6 (i.e. reference frame), i.e. those section for which changes were visible. B Overlay of protomers in L, T, O of (1) AcrBZ crystal structure (PDB-ID: 5NC5) (in grey; partially transparent) and (2) cryo-EM derived AcrBZ structure reconstituted in *E. coli* lipids inside a saposin A-nanodisc. The overlay was produced using MatchMaker command in Chimera using the rigid TMH 4-6 (blue) of AcrB crystal structure as reference frame.

In the AcrBZ structure in saposin A discs, we noted a previously unseen, significant bending of AcrZ towards the binding groove in AcrB, which was most pronounced in L and O states. Each of the three AcrZs in the saposin disc bends towards AcrB between residues 10 and 15 (F10, A11, V12, I13, M14, V15) (Fig 4A). The bending mode is also predicted by molecular dynamics simulations of the AcrBZ complex in a lipid bilayer (Fig 4B). The bend could be caused by the presence of lipids in the saposin disc mediating new contacts with AcrB or otherwise exerting a force on AcrZ.

**Figure 4.**
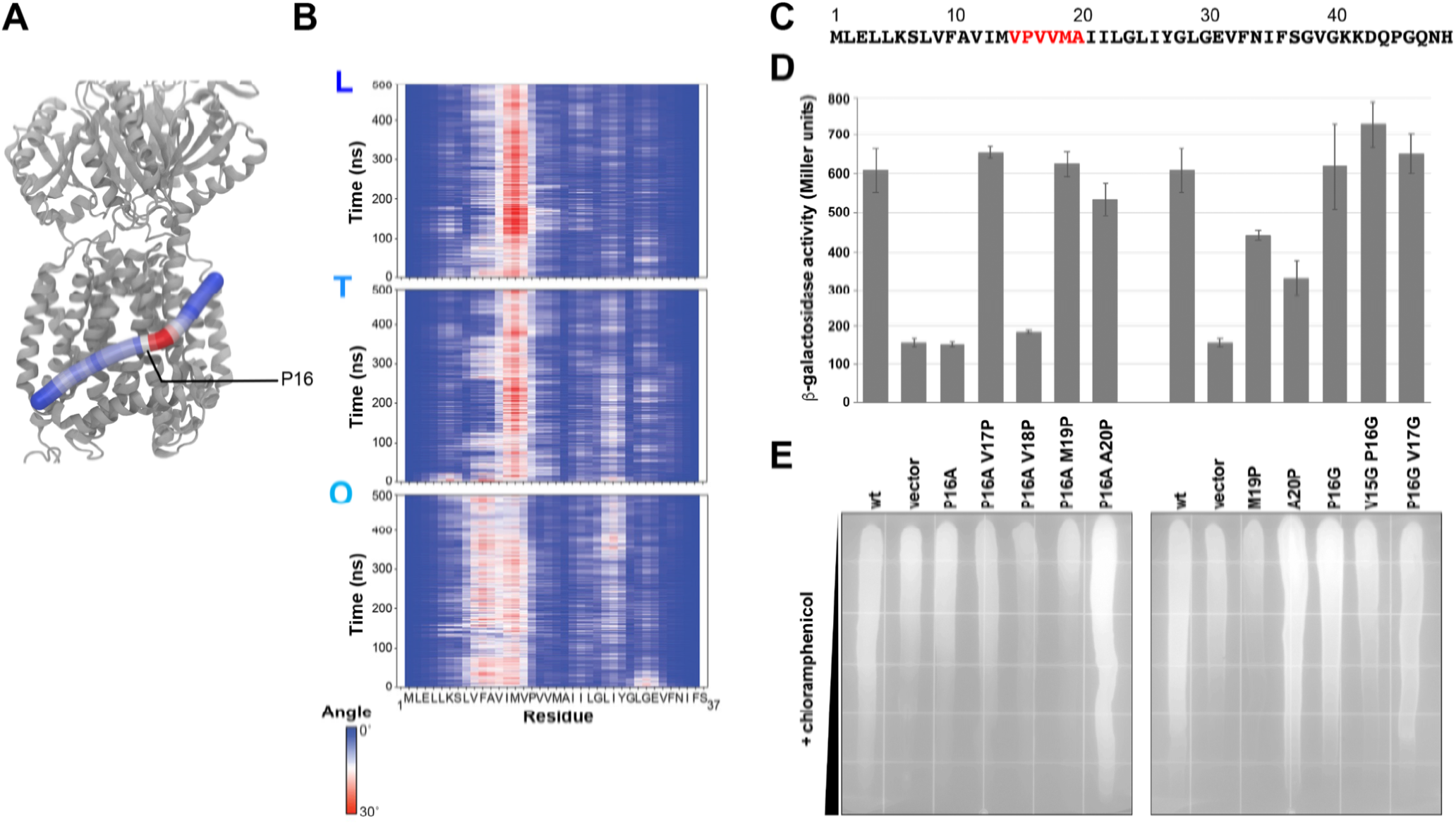
Impact of AcrZ curvature on binding to AcrB. A A snapshot of AcrZ bound to the AcrB in the loose conformation. The AcrZ is shown in tube representation and coloured based on the bending angle between its residues, from blue (0°) to red (30°), while the AcrB is shown in ribbon representation and coloured grey. B The evolution of this bending angle over the course of one of the simulations for all three AcrZ subunits. The conformation of AcrB (L, T, O) to which these AcrZ subunits are bound is shown on top left of each graph. C The sequence of AcrZ with residues mutated indicated in red. D Split adenylate cyclase two-hybrid assays of the interaction between plasmid-encoded T25-AcrB and the empty vector, wild type AcrZ-T18 or the indicated mutant. T25-AcrB and the AcrZ-T18 indicated were co-expressed in an adenylate cyclase deficient strain and grown to OD_600_ ∼ 1 when cells were harvested for β-galactosidase activity assay. Shown are the average and standard deviation of three experiments. The wt and vector samples are each shown twice. E Exponentially-growing cultures of the *E. coli* Δ*acrZ* strains carrying the pBAD24 empty vector, wild type AcrZ or the indicated AcrZ mutant were applied across chloramphenicol gradient plates to visualize differences in antibiotic sensitivity. The plates were incubated overnight at 37°C and photographed. Shown here is a representative image of an experiment carried out in triplicate.

### Defined bend in AcrZ important for interaction with AcrB and effect on efflux

A proline at position 16 is expected to be a helix-breaker (Fig 4C), conferring a kink to the AcrZ protein due to the R-group ring. Given the significant bending seen in this region in the structural analysis, we asked whether the proline residue is important for the AcrZ interaction with AcrB. Using a two-hybrid approach with AcrB and AcrZ fused to the T25 and T18 fragments of *Bordetella pertussis* adenylate cyclase, respectively, we assayed for interactions that restore adenylate cyclase function, and consequently induce β-galactosidase activity (Hobbs *et al*, 2012). We first substituted an alanine for the proline residue (P16A) and observed that for this mutant the β-galactosidase activity was equivalent to empty vector levels, indicating the P16A variant could not interact with AcrB (Fig 4D). We then shifted the location of the proline from position 16 to positions 17, 18, 19 and 20 (P16A V17P, P16A V18P, P16A M19P, P16A A20P). With the exception of P16A V18P, all mutations restored β-galactosidase activity to wild type levels (Fig 4D). We next examined the consequences of introducing a second proline at positions 19 and 20 (M19P, A20P). The M19P and A20P variants had reduced β-galactosidase compared to wild type. Finally, we investigated the effect of changing flexibility in this region by introducing single and double glycine substitutions (P16G, V15G P16G, P16G V17G). In principle, glycine could permit the AcrZ to bend in the same conformation favoured by proline, and indeed all glycine substitution AcrZ variants were able to interact with AcrB at wild type levels (Fig 4D). Together, these results indicate that a bend in AcrZ around position 16 is important for the AcrZ-AcrB interaction.

To investigate the effects of AcrZ mutations on drug efflux, untagged derivatives were expressed from a plasmid in a Δ*acrZ* strain background and assayed for resistance to chloramphenicol. Compared to the vector control, wild type *acrZ* provided increased resistance to chloramphenicol on gradient plates (Fig 4E). As expected, mutant AcrZ variants unable to interact with AcrB were unable to rescue chloramphenicol resistance. However, several AcrZ mutants capable of interacting with AcrB were nonetheless unable to restore growth to wild type levels (V15G P16G; P16G V17G). Interestingly two mutants that had a wild type or intermediate phenotype for AcrB interaction (P16A M19P; M19P) showed reduced chloramphenicol resistance to below that of the empty vector, while two other mutants with a wild type AcrB interaction (P16A A20P; A20P) were more resistant than the wild type AcrZ strain.

Interestingly, mutations of individual interfacial residues to alanine did not have a strong effect on either the AcrZ interaction with AcrB or chloramphenicol resistance (Appendix Fig S3). Together, these results indicate that the overall hydrophobic character and bent shape of AcrZ, rather than specific AcrB-AcrZ contacts, are important for physical interactions. Moreover, the mutations also indicate that direct interactions of AcrB-AcrZ are necessary but not sufficient for efflux function, and that the interactions can either suppress or support pump activity depending on context. These features are a hallmark of an allosteric system.

### Lipid interactions with AcrB and AcrBZ

It had previously been suggested that lipids can modulate the structure and function of specific membrane proteins, and cardiolipin has been identified as such a modulator (Bechara *et al*, 2015; Gupta *et al*, 2017; Laganowsky *et al*, 2014). We carried out coarse-grained molecular dynamics simulations of AcrB and AcrBZ with cardiolipin and palmityloleoyl phosphatidylglycerol (POPG), both of which are the less abundant lipids found in the *E. coli* cytoplasmic membrane. Our simulations started with no cardiolipin and POPG molecules within 30 Å of the protein, but after 5 µs as many as 15 cardiolipin and 50 POPG molecules were found within 6 Å of the protein (Appendix Fig S4). The cardiolipin and POPG enriched around both proteins despite the membrane containing a smaller number of these lipid types compared to palmityloleoyl phosphatidylethanolamine (POPE). A slightly higher degree of enrichment was observed around AcrBZ compared to AcrB, especially in the case of POPG. Analysis of density plots suggest that the interaction between these lipids with AcrB or AcrBZ may be non-specific, as high-density regions were found around all parts of the transmembrane portion of the proteins (Appendix Fig S4). The higher number of cardiolipin and POPG is likely to result from the increase protein surface area (Appendix Fig S4D). Given that it is easier to genetically manipulate the cardiolipin composition of the *E. coli* inner membrane, we focussed our functional and structural studies on this special lipid.

The *E. coli* inner membrane has been reported to contain around 5% cardiolipin (Dowhan, 1997), with some localized enrichment in negatively curved regions of the membrane such as the cell poles (Renner & Weibel, 2011). To test the effects of cardiolipin on the conformation of AcrB and AcrBZ, we reconstituted the proteins into saposin discs in which the *E. coli* lipids were supplemented with an additional 5% cardiolipin (i.e., to ∼10% total abundance). The increase in the cardiolipin content is associated with small changes in PC1/2 and PN2 for AcrB (Appendix Fig S2). Minor changes were seen when comparing the AcrBZ complexes in the presence and absence of excess cardiolipin. The greatest changes were seen in comparing AcrB with natural lipids with AcrBZ in the lipid environment with added cardiolipin (Appendix Fig. S2).

### Additive effects of AcrZ and cardiolipin for growth viability and drug binding

*E. coli* lacking AcrZ was previously shown to be less resistant to chloramphenicol compared to the wild type parental strain (Hobbs *et al*., 2012). Due to the conformational changes in AcrB observed in the presence of AcrZ and additional cardiolipin, we asked whether cardiolipin could affect chloramphenicol resistance. Thus the wild type strain (*E. coli* MG1655) and strains lacking AcrZ (MG1655 Δ*acrZ*), cardiolipin (MG1655 Δ*clsABC*::FRT-*kan*-FRT) or both (MG1655 Δ*clsABC*::FRT-*kan*-FRT Δ*acrZ*) were assayed for chloramphenicol resistance. As reported previously, the *acrZ* deletion strain showed increased sensitivity to chloramphenicol but not erythromycin (Hobbs *et al*, 2012) (Fig 5). Strains deficient for cardiolipin also were more sensitive to chloramphenicol. Strikingly, the double mutant strain was most sensitive to chloramphenicol, consistent with the largest structural changes being seen in the presence of both AcrZ and cardiolipin. These results indicate AcrZ and cardiolipin have additive effects on AcrB ability to export chloramphenicol (Fig 5).

**Figure 5.**
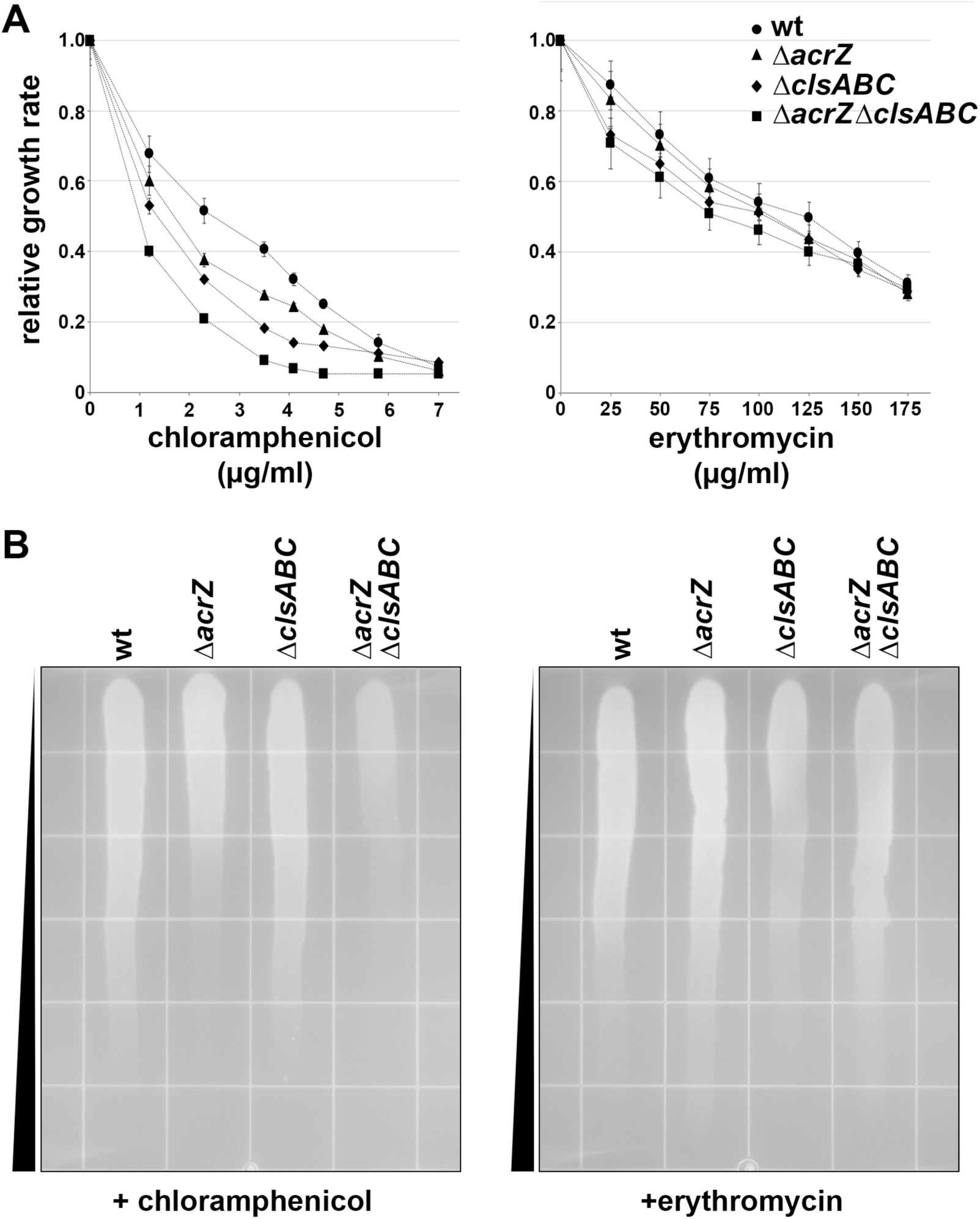
Cardiolipin biogenesis and AcrZ impact on chloramphenicol sensitivity. A Growth rates of *E. coli* MG1655 (wild type parent strain), MG1655 Δ*acrZ*, MG1655 Δ*clsABC*::FRT-*kan*-FRT (cardiolipin-deficient) and MG1655 Δ*clsABC*::FRT-*kan*-FRT Δ*acrZ* (cardiolipin-deficient and Δ*acrZ*) in the presence of a range of chloramphenicol (0 to 7 µg ml^-1^) and erythromycin (0 to 175 µg ml^-1^) concentrations were determined relative to the maximum growth rate in the absence of drug. OD_660_ measurements are presented as mean of triplicate measurements ± standard error of the mean. For the determination of the relative growth rates of the cultures in each of the wells, the exponential phase of the growth curve (as a mean of n = 3 for each culture type) was determined from the linear increase in a log_10_(OD_660_) versus time plot. The slope of this section was determined by simple linear regression. Heteroscedasticity-consistent standard errors of the corresponding slope coefficient were calculated. The quality of the fit was significant in all cases (P < 0.05). Next, the relative growth rate was determined as the ratio of the growth rate in the presence of drug over the maximum growth rate in the absence of drug. B Exponentially-growing cultures of the above strains were applied across chloramphenicol or erythromycin gradient plates to visualize differences in antibiotic sensitivity. The plates were incubated overnight at 37°C and photographed. Shown here is a representative image of an experiment carried out in triplicate.

### Combination of AcrZ and cardiolipin is associated with changes in the AcrB entry channels, gating loop and binding pocket

Efflux of substrates through AcrB is complex process given that AcrB has multiple entry channels and multiple binding sites that are used by different types of drugs (reviewed in (Zwama & Yamaguchi, 2018). Deletion of *acrZ* affects only a subset of the antibiotics effluxed by AcrB. The mechanism behind this specificity is unknown, but it is possible that AcrZ is selectively affects certain entry channels or binding locations. Thus, we examined structural differences in AcrB regions involved in drug transport in the presence and absence of AcrZ and cardiolipin. We found AcrZ and cardiolipin affect both of the substrate entry channels (Fig 6A and Appendix Fig S5). Channel 1, for entry from the periplasm, has an altered entry shape with closer access from above the membrane surface in AcrBZ compared to AcrB. Channel 2, which protrudes sideward from above the outer leaflet of the membrane into the protomer in the L state, is more restricted by a loop region part of PC1/2 of the AcrB structure in a saposin disc without cardiolipin enrichment when compared to AcrBZ structure in the saposin disc with ∼10% cardiolipin.

**Figure 6.**
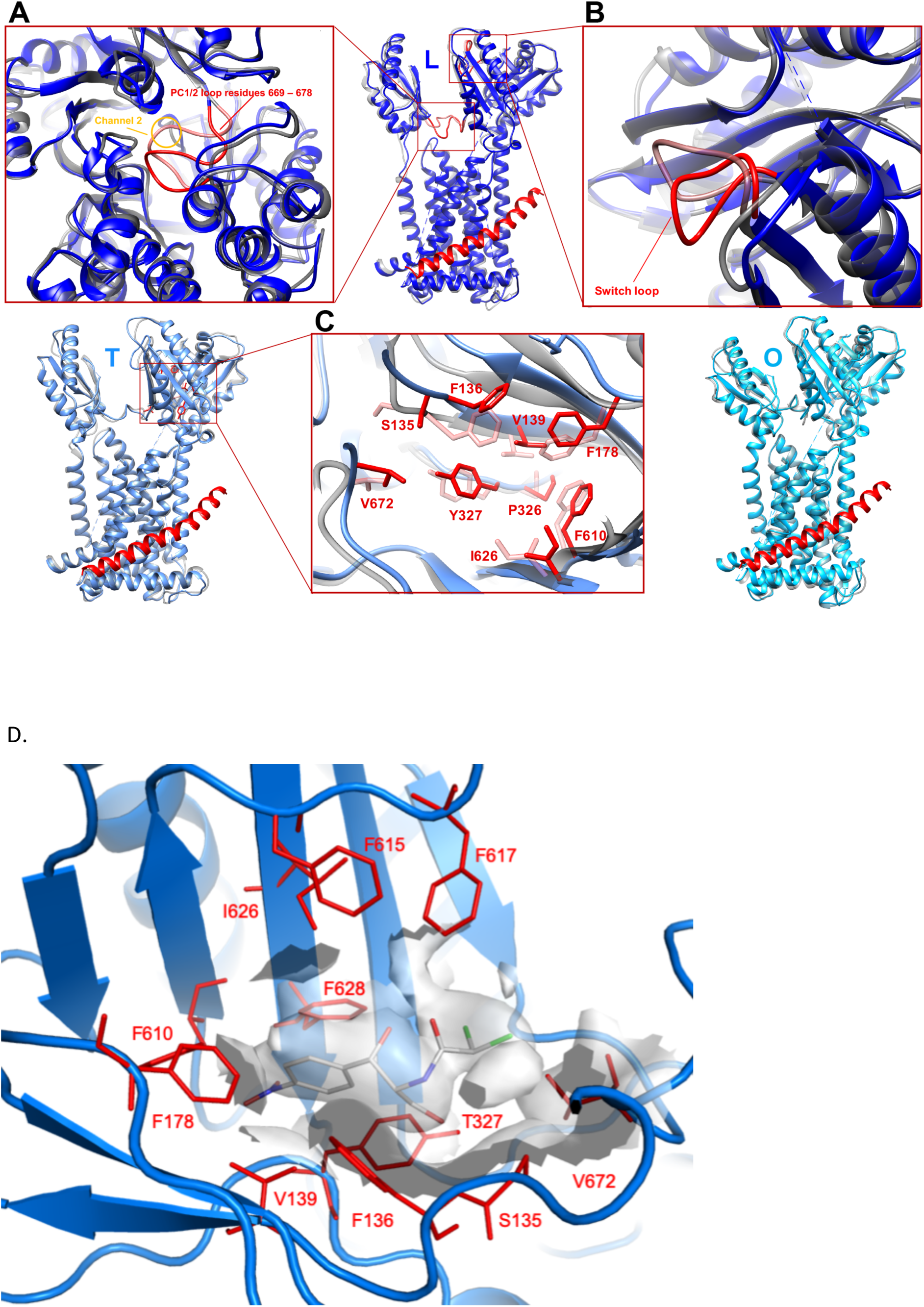
Structural comparison between saposin A-disc reconstituted AcrB and AcrBZ with cardiolipin enrichment. Overlay of protomers in L, T, O of cryo-EM derived AcrB (in grey; partially transparent) reconstituted in *E. coli* lipids inside a saposin A-disc and AcrBZ (blue variants) reconstituted in *E. coli* lipids enriched with cardiolipin inside a saposin A-disc. A Channel 2 entry is restricted by a loop region of PC1/2 (red) for substrate entry from the outer leaflet of the inner membrane in case of AcrB in L state but open in the case of the AcrBZ complex in L. B Once substrate enters the protomer in L, a switch loop (red) allows passage into the deep-binding pocket in AcrB in complex with AcrZ. This loop appears to restrict access in case of the AcrB only protomer. C Impact on the drug binding pocket at the site of chloramphenicol binding. Chloramphenicol (not depicted here) is located inside the distal pocket of the AcrB in ‘tight’ conformation. The density here is not sufficiently well resolved to precisely model the detailed contacts, but the antibiotic is likely to pack against residues P326, Y327, V139, F136, F610, F178, S135, I626, and V672. The residues are mostly on two sets of beta sheets in the porter domain of AcrB, as well as on nearby loops. D CryoEM density in the T state pocket for the AcrBZ complex with additional cardiolipin. The density shape does not uniquely define the orientation of the chloramphenicol, and the model shows one possible conformation. Density was not observed in this position for the maps for the AcrBZ and AcrB structures in the natural lipid composition or AcrB with cardiolipin supplement.

There are also changes in the switch loop that could influence the passage of the drug from the L to the T states (Fig 6B). To explore the process of substrate movement through the pocket, molecular dynamics simulations were conducted in which a chloramphenicol molecule was pulled from the periplasmic space into the deep binding pocket of the L protomer of AcrB and AcrBZ with 10% cardiolipin. However, the results suggest that the movement of the antibiotic is unaffected by the switch loop at the entry gate of the ligand from the L to the T state due to its inherent flexibility (Appendix Fig S6).

The observed changes in PC1/2 and PN2 of AcrB are likely to affect drug-binding pocket properties in the T protomer given the orientation of key residues for substrate binding (Fig 6C). When AcrBZ saposin discs with supplementary cardiolipin were prepared with the addition of chloramphenicol, additional density was found in the cryo-EM map in the distal pocket of the AcrB protomer in the T state, which could correspond to the antibiotic bound in a number of discrete conformations, averaged together in the process of EM image reconstruction (Fig 6D). There was no apparent density for chloramphenicol in the corresponding pocket when the disc was not supplemented with cardiolipin or when AcrZ was not present. Molecular dynamics simulations also indicate that chloramphenicol is bound more stably to the binding site of the T protomer of the AcrBZ + 10% cardiolipin structure compared to the AcrB structure (Appendix Fig S7). Chloramphenicol significantly and frequently changed its orientation and failed to reach a stable conformation in the simulations of AcrB. These data suggest that the presence of AcrZ and cardiolipin can influence the occupancy of the substrate in the binding pocket. Together the structures and simulations show that, although the effects of AcrZ and cardiolipin on the transport activity of AcrB might not occur at the substrate encounter with the switch loop, multiple other steps are affected.

## Discussion

The development of a procedure for efficient reconstitution of AcrBZ into saposin A-scaffold based discs with a native lipid environment, suitable for cryo-EM analysis and with an option to alter the lipid content, revealed previously undescribed structural differences in the conformation of AcrZ. The structure, together with mutagenesis data, revealed that the overall bent shape of AcrZ is being recognized rather than specific interacting residues, which also may be the case for the transmembrane interactions of other small proteins with their cognate partners (Hobson *et al*, 2018). The elucidation of a more native conformation of both AcrB and the AcrBZ complex helps to explain the mechanism behind the resistance-modulating effect of AcrZ binding to AcrB for a subset of its substrates.

In light of the discovery of AcrZ, there appears to be a crucial yet unexplored general role of small proteins for modulating activity of efflux pumps. For example, in muscle cells, a group of small proteins regulate calcium uptake by the SERCA pump (Anderson *et al*, 2015). In *E. coli*, KdpF is another small protein that increases the activity of a potassium transport protein (KdpFABC), while SgrT is a small protein that inhibits the glucose transport activity of EIICB (Storz *et al*, 2014). Structural studies of the role of these small proteins can further elucidate the different mechanisms of possible allosteric modulation. A number of other small proteins detected in *E. coli* are also found at the membrane (Hemm *et al*, 2008), and we speculate that many of these proteins can act as allosteric modulators of target membrane protein partners.

Accumulating evidence indicates that lipids can affect the localization, structure, stability and function of certain membrane proteins (Bechara *et al*, 2015; Gupta *et al*, 2017; Laganowsky *et al*, 2014) (Renner & Weibel, 2011). Lipids have, for instance, been shown to directly regulate opening of the mechanosensitive channel MscS (Pliotas *et al*, 2015). *E. coli* aquaporin Z was found to be stabilised and functionally modulated by cardiolipin, and the *E. coli* ammonia channel AmtB was shown to selectivity bind phosphatidylglycerol (Laganowsky *et al*, 2014). Another study with co-polymer solubilized AcrB identified a pattern of lipid organization around the transmembrane portion (Qiu *et al*, 2018). We observe lipids in our maps as well, and our best model for the AcrBZ complex with lipids is consistent with acyl chain packing and lipid head group interactions with AcrB observed in the other study (Fig 2).

We structurally, computationally and functionally explored the impact of cardiolipin on AcrBZ. Taken together, our data suggest that the interaction of AcrZ and AcrB increases affinity of the complex for cardiolipin- and POPG-enriched environments inside the inner membrane. This effect does not appear to be due to the formation of specific interactions, but instead may originate from the greater surface for lipid contact in the presence of the AcrZ subunit (Appendix Fig S4). Cardiolipin and other lipids, together with AcrZ, modulate the activity of AcrB through allosteric changes – putatively by inducing structural alterations in the drug entry and binding sites. The activities of other bacterial and mitochondrial membrane proteins have also been reported to be affected by interactions with cardiolipin (Dudek, 2017). Both a small protein and cardiolipin have been separately shown to promote the activity of MgtA, a magnesium importer in *E. coli* (Subramani *et al*, 2016; Wang *et al*, 2017a). Cardiolipin thus impacts processes ranging from electron transport to antimicrobial resistance by affecting protein localization, enhancing protein stability, mediating interactions between monomer units, and transmitting conformation changes between subunits (Dudek, 2017). Additionally, recent work has found that cardiolipin plays a role in proton motif force stimulation and modulation of ATPase activity in SecYEG (Corey *et al*, 2018). Cardiolipin promotes the distribution of the osmosensory transporter ProP to the cellular poles in *E. coli* (Romantsov *et al*, 2007). Interestingly, in older bacterial cells, the AcrAB/TolC complex tends to cluster at the pole (Bergmiller *et al*, 2017), where it potentially could encounter a distinct lipid environment with impact on its function and activity. The recent studies of protein-lipid interactions, and cardiolipin in particular, shows the importance of continued study of membrane proteins in their native lipid environment. Knowledge of these interactions may impact on understanding the mechanism of drug resistance in clinical treatment.

## Materials and Methods

### Strains, plasmids, primers

Constructs of AcrB/DARPin and AcrBZ/DARPin complexes were previously described (Du *et al*, 2014; Eicher *et al*, 2012). Strain MG1655 Δ*acrZ*::*km* also was described earlier (Hobbs *et al*, 2012). Strain MG1655 Δ*clsABC* was kindly provided by Douglas B. Weibel (Oliver *et al*, 2014). To make the Δ*acrZ* Δ*clsABC* double mutation strain, the Δ*acrZ*::*km* allele was transduced from GSO213 (Hobbs *et al*, 2012) into MG1655 Δ*clsABC* strain following a standard P1 transduction protocol for *E. coli* genome manipulation (Thomason *et al*, 2007). MG1655 Δ*acrZ* (GSO284) was the background strain for all assays of *acrZ* mutants (Hobbs *et al*, 2010). BTH101 was the background strain for the two-hybrid assays (Hobbs *et al*, 2012).

### Preparation of saposin A

Human saposin A was expressed in *E. coli* shuffle-T7 cells. Colonies of shuffle-T7/pET15b-saposin A from a freshly transformed plate were used to inoculate 50 ml of LB medium containing carbenicillin (100 μg/ml) in a 250-ml baffled flask. The cells were grown in an orbital shaker at 30°C, 220 rpm overnight. 20 ml of the starter culture was used to inoculate 1000 ml LB medium (with carbenicillin) in a 2-l baffled flask at 30°C, 220 rpm, and cultures were induced at A_600_=0.6 – 0.8 with 1 mM isopropyl 1-thio-b-D-galactopyranoside (IPTG). The temperature was dropped to 16°C and the cells were grown at 220-rpm overnight. Cells were harvested by centrifugation at 4,200 rpm for 25 min at 4°C. Cell pellets from a 2-l culture were resuspended in 50 ml AEX buffer (50 mM Tris-HCl, pH 7.4, 25 mM NaCl) with 1 tablet EDTA free protease inhibitor cocktail tablets and 5U/ml DNase I. Cells were lysed using a homogenizer (Emulsiflex) at 15,000 psi and the cell debris was pelleted by centrifugation at 40,000 xg for 30 min at 4°C. The supernatant was collected in a glass bottle, and was heat-treated at 85°C in a water bath for 10 min with gentle shaking, followed by centrifugation at 40,000 xg for 30 mins at 4°C. The supernatant was loaded onto a HiTrap Q column (GE Healthcare) equilibrated with AEX buffer. The column was washed with 10 column volumes of AEX buffer and then saposin A was eluted over a 0 – 50% gradient of AEX elution buffer (20 mM Tris, pH 7.4, 1 M NaCl). The fractions containing saposin A were pooled, and the protein was concentrated to 0.5 ml using 3 kDa Vivaspin concentrator. A final gel filtration step was performed using a Superdex 200 column equilibrated with GF Buffer-1 (20 mM HEPES pH 7.0, 150 mM NaCl). The peak fractions were concentrated, flash frozen in liquid nitrogen and stored at −80°C.

### Protein expression, purification and nanodisc reconstitution

AcrB/DARPin and AcrBZ/DARPin complex purification followed the procedure described earlier (Du *et al*, 2014). The purified proteins were concentrated, frozen in liquid nitrogen and stored at −80°C.

Nanodiscs were reconstituted using a modification of a procedure described earlier (Frauenfeld *et al*, 2016). Efficient nanodisc formation requires a step at pH 4, but AcrB and AcrBZ were unstable under this acidic condition. Thus, in the first step of the reconstitution procedures, nanodiscs were prepared with *E. coli* lipids at acidic pH and then brought to pH 7. In the second step, purified AcrB and AcrBZ were reconstituted into the pre-formed nanodiscs at the neutral pH.

For lipid stock preparation, 20 mg of *E. coli* total lipid extract (Avanti) was dissolved in 0.5 ml of chloroform. For lipid stock with extra cardiolipin, 20 mg of *E. coli* total lipid extract and 1 ml of cardiolipin were mixed and dissolved in 0.5 ml of chloroform. The lipid solutions in glass vials were evaporated in a vacuum desiccator, then resuspended in 1 ml of 50 mM HEPES pH 7.5, 150 mM NaCl and sonicated for 30 min. The lipid stocks were stored at −20°C. For the nanodisc reconstitution, 28.2 µl of saposin A (4.484 mM) was mixed with 50.8 µl of *E. coli* lipid stock (25 mM; for cardiolipin-enriched nanodiscs, 5% v/v purified cardiolipin (Avanti) was added to the lipid stock used for nanodisc formation), and then sodium acetate (50 mM pH:4.8) was added to the mixture to a final volume of 500 µl and incubated at 37°C for 10 min. 1 ml of GF buffer-2 (Tris 20 mM pH 7.5, NaCl 150 mM) was added to the mixture, and the buffer was exchanged to GF Buffer-2 using a HiTrap Desalting 5 ml column. 2 ml of the eluate was mixed with AcrB/DARPin or AcrBZ/DARPin. The molar ratio of AcrB/DARPin or AcrBZ/DARPin:saposin A:lipid is about 1:10:100 in the reconstitution. The mixture was incubated at 4°C for half an h and dialyzed against 1000 ml of GF Buffer-2 overnight. The sample was dialyzed again against 1000 ml of GF Buffer-2 for further 3 h. The buffer exchanged sample was concentrated to 500 µl and purified by gel filtration chromatography using a Superdex 200 column equilibrated with GF Buffer-2. The peak fractions containing the protein in nanodiscs were concentrated to 2 mg/ml. Minocycline was added to AcrB/nanodisc or AcrB/nanodisc-cardiolipin to a final concentration of 2 mM; Chloramphenicol was added to AcrBZ/nanodisc or AcrBZ/nanodisc-cardiolipin to 1 mM. The mixtures were incubated for 1 h at 4°C, then flash frozen in liquid nitrogen and stored at −80°C.

### Cell growth for drug sensitivity assay

Colonies from overnight cultures of freshly transformed and streaked out plates of *E. coli* MG1655 (wild type parent strain), MG1655 Δ*acrZ*, MG1655 Δ*clsABC*::FRT-*kan*-FRT (cardiolipin-deficient) and MG1655 Δ*clsABC*::FRT-*kan*-FRT Δ*acrZ* (cardiolipin-deficient and Δ*acrZ*), were picked and streaked out again onto sterile LB-Agar plates and grown overnight at 37°C. These colonies were picked and grown in LB for about 3 h at 37°C, and then used to inoculate fresh medium in a 96-well plate. The cells were diluted to OD_600nm_ ∼0.06 in fresh LB medium in the wells of a 96-well plate to which antibiotics were added as indicated in Fig 5. Growth was followed over time at OD_660_ at 37 °C in a CLARIOstar microplate reader (BMG LABTECH). For the determination of the relative growth rates of the cultures in each of the wells, the exponential phase of the growth curve (as a mean of n = 3 for each culture type) was determined from the linear increase in a log_10_(OD_660nm_) versus time plot. The slope of this section was determined by simple linear regression. Heteroscedasticity-consistent standard errors of the corresponding slope coefficient were calculated. The quality of the fit was significant in all cases (P < 0.05). Next, the relative growth rate was determined as the ratio of the growth rate in the presence of drug over the maximum growth rate in the absence of drug.

### Electron cryo-microscopy

For the structure determination of saposin A discs with AcrB and AcrBZ as well as both complexes with supplementary cardiolipin, a 4.0 µl aliquot at 2 mg/ml was applied onto holey carbon film supported by a 300-mesh R1.2/1.3 Quantifoil gold grid (Quantifoil) that had been previously glow discharged. The grid was blotted for 2.5 – 3.0 seconds and rapidly frozen in liquid ethane using a Vitrobot IV (FEI) at 4°C and 100% humidity. The grids were stored in liquid nitrogen before imaging. Zero-energy-loss images of frozen-hydrated AcrB disc particles were recorded automatically on an FEI Titan Krios electron microscope at 300 kV, using a slit width of 20 eV on a GIF Quantum energy filter and a Gatan K2-Summit direct electron detector (Gatan) in counting mode. Images of frozen-hydrated AcrB/nanodisc-cardiolipin, AcrBZ/nanodisc and AcrBZ/nanodisc-cardiolipin particles were acquired automatically on the FEI Titan Krios electron microscope at 300 kV using a Falcon III direct electron detector camera (FEI) in counting mode.

### Image processing and 3D reconstruction

The software MotionCor2 (Zheng *et al*, 2017) was used for whole-frame motion correction and dose weighting, Gctf (Zhang, 2016) for estimation of the contrast transfer function parameters, and RELION-3.0 (Scheres, 2012) package for all other image processing steps. A particle subset was manually selected to calculate reference-free 2D class averages, which was then used as templates for automated particle picking of the entire data set. The templates were low-pass filtered to 20 Å to limit model bias. Then several runs of 2D classifications were used to remove the heterogeneous particles, as well as the false positive particles from the auto-picking. A reference map was generated from crystal structure of AcrB using a program pdb2mrc in the EMAN package (Ludtke *et al*, 1999), and was low pass filtered to 60 Å resolution and was used as a starting point for the 3D classification. We selected good particles for further analysis based on the quality and high resolution in the 2D and 3D classification. The 3D auto-refinement resulted in near-atomic resolution maps. After per-particle motion correction and radiation-damage weighting by Bayesian polishing in RELION (Zivanov *et al*, 2018), the polished particles were subjected to 2D and 3D classifications, and 3D auto-refinement again. A soft mask in RELION post-processing was applied before computing the FSCs. The overall resolution of the maps was estimated by the gold-standard FSC criterion with 0.143 cut-off (Henderson and Rosenthal, 2003). Local resolution variations were estimated with ResMap (Kucukelbir *et al*, 2014). The data collection and processing parameters for all four specimens are summarized in Table S1.

### Model docking and refinement

Automated structural refinement using Rosetta followed the procedure described in (Wang *et al*, 2017b). The models were refined with PHENIX (Adams *et al*, 2010) and ISOLDE (Croll, 2018), and the structures were further modelled and visualized using PyMOL (DeLano, 2002), Chimera (Pettersen *et al*, 2004) and Coot (Emsley *et al*, 2010). The crystal structure of human saposin A in the open state was docked into the maps. Density was also apparent for lipids, and the corresponding hydrocarbon portion was modelled into the density where there was good correspondence. Although the AcrB/nanodisc specimens were prepared in the presence of minocycline, density for the compound was not apparent in the refined cryo-EM maps inside the periplasmic domain. However, weak density that is consistent with minocycline is present near F556 (subunit A). A more defined density was found in the AcrBZ/nanodisc-cardiolipin EM map and subsequently assigned to chloramphenicol, which was added to the sample prior to freezing. The quality of the stereochemistry was evaluated with EMRinger (Barad *et al*, 2015) and MOLPROBITY (Chen *et al*, 2010).

### Molecular dynamics simulations

For the atomistic simulations, the proteins were parameterized using CHARMM36 force field (Huang & MacKerell Jr, 2013), whilst the parameters for the antibiotic chloramphenicol were obtained using CHARMM-GUI ligand modeler (Kim *et al*, 2017). The AcrZ P16A M19P mutant was generated using PyMOL (DeLano, 2002). A small patch of 15 × 15 nm model membrane was constructed using CHARMM-GUI membrane builder (Jo *et al*, 2009) to mimic the lipid composition of *Escherichia coli* K12 inner membrane (Aibara *et al*, 1972; Lugtenberg & Peters, 1976; Yokota *et al*, 1980) (75% 1-palmitoyl 2-cis-vaccenic phosphatidylethanolamine (PVPE), 20% 1-palmitoyl 2-cis-vaccenic phosphatidylglycerol (PVPG) and 5% 1-palmitoyl 2-cis-vaccenic 3-palmitoyl 4-cis-vaccenic diphosphatidylglycerol (cardiolipin)). Protein insertion into the membrane was performed using the *g_membed* protocol in GROMACS (Wolf *et al*, 2010). This system was then solvated with the SPC water molecules (Lins & Hünenberger, 2005) and neutralised with 0.15 M NaCl. A short 1 ns equilibration simulation was performed whereby the heavy atoms of the protein were positionally restrained using a force constant of 1000 kJ mol^-1^. The temperature was kept at 310 K using the Nose-Hoover thermostat with a time constant of 1.0 ps (Hoover, 1985; Nosé, 1984). The pressure was held at 1 atm using a semi-isotropic coupling to the Parrinello-Rahman barostat with a time constant of 1.0 ps (Parrinello & Rahman, 1981b). The electrostatic interactions were calculated using the smooth particle mesh Ewald method with a real-space cut off value of 1.2 nm (Essmann *et al*, 1995). The twin range cut-off method with a cut-off value of 1.2 nm was used to calculate the Van der Waals interactions, whereby the forces were switched to zero between 1.0 to 1.2 nm. The LINCS algorithm was utilised to constrain all covalent bonds to their equilibrium value, allowing for a 2 fs time step (Hess *et al*, 1997). After this equilibration simulation, the position restrains on the protein were removed and two independent production runs, each for 500 ns, were conducted with different starting velocities. Steered molecular dynamic simulations were performed whereby a harmonic spring with a force constant of 100 kJ mol^-1^ nm^-2^ was attached to the center of mass of chloramphenicol and pulled at a constant velocity of 0.1 nm ns^-1^ towards a reference residue (F136) found in the deep binding pocket of the L protomer. Three independent steered molecular dynamic simulations were performed for each of AcrB and AcrBZ starting with different velocities.

For the coarse-grained simulations, the proteins were converted to coarse-grained representation using the *martinize.py* script using the MARTINI 2.2 force field (Monticelli *et al*, 2008) with ElNeDyn to retain the secondary and tertiary structures (Periole *et al*, 2009). A patch of 30 x 30 nm membrane model of the same lipid composition as the atomistic simulation was constructed using CHARMM-GUI Martini Maker Bilayer Builder (Qi *et al*, 2015). The protein was placed in the middle of the membrane and overlapping lipids were removed. To understand the interactions between the protein and charged lipids, we reorganized the position of POPG and cardiolipin such that at the beginning of the simulation there were none of these lipids within 30 Å of the protein. The simulation box was solvated with the standard MARTINI water molecules and neutralised with 0.15 M NaCl. Energy minimisation was then performed using the steepest descent method. The system was subsequently equilibrated for 10 ns with positional restrains applied on the protein. The temperature was maintained at 310 K using a velocity-rescaling thermostat (Bussi *et al*, 2007), with a relaxation time of 1 ps, whereas the pressure was maintained at 1 bar by a semi-isotropic coupling with the Berendsen barostat (Berendsen *et al*, 1984) and a time constant of 5 ps. The cut-off for the nonbonded interactions was set at a distance of 1.2 nm, while the Lennard-Jones and Coulomb potentials were shifted from 0.9 and 0.0 to the cut-off distance, respectively. The LINCS algorithm was used to constrain all covalent bonds to their equilibrium values (Hess *et al*, 1997) and the time step was slightly increased from 2, 5, 8 to 10 fs to allow the system to be well equilibrated. After the equilibration simulations, protein position restraints were removed and three independent production runs with different starting velocities were performed for 5 μs using the same parameters, except a Parrinello-Rahman barostat with a time constant of 12 ps was used to control the pressure (Parrinello & Rahman, 1981a).

All simulations were performed using the GROMACS 5 package (Abraham *et al*, 2015) and visualised in VMD (Humphrey *et al*, 1996). The bending angle of AcrZ was determined using the Bendix plug-in within VMD (Dahl *et al*, 2012). Protein-lipid contact analysis was performed using *gmx select*. The partial mass density landscapes was generated with a modified version of the *g_density* tool (Castillo *et al*, 2013).

### Bacterial two-hybrid assays

Single colonies of a *E. coli* strain adenylate cyclase deletion strain (BTH101) freshly transformed with plasmids bearing *acrZ* fused to the T18 fragment of adenylate cyclase on the C-terminus and *acrB* fused to the T25 fragment of adenylate cyclase on the N-terminus were grown in 1 ml of LB medium supplemented with 30 µg/ml kanamycin and 50 µg/ml carbenicillin at 37°C with shaking at 250 rpm overnight. The cultures were diluted 1:100 into 3 ml of the same medium also containing 1 mM IPTG inducer and were similarly grown to OD_600_ ∼ 1. An aliquot of sample was used for β-galactosidase assays as previously described (Miller, 1992) with some modifications. The OD_600_ was recorded and a 100 µl aliquot was added to a 1.5 ml microfuge tube containing 700 µl Z buffer (60 mM Na_2_HPO_4_, 40 mM NaH_2_PO_4_, 10 mM KCl, 1 mM MgSO_4_, 2.7 µl/ml of β-mercapthoethanol, 1.5 µl/ml of 0.1% SDS) and 30 µl chloroform. The samples were vortexed and incubated for 15 min at 28°C, whereupon 100 µl of 8 mg/ml ortho-nitrophenyl-β-galactoside (ONPG) in Z buffer was added to the sample and vortexed and returned to 28°C. Once a yellow colour developed (∼10 min), 0.5 ml of 1 M Na_2_CO_3_ was added and the sample was vortexed. The start times (addition of ONPG) and the stop times (addition of Na_2_CO_3_) were recorded. The sample was centrifuged to remove pellet debris and the A_420_ of the supernatant was recorded. Miller Units were calculated as follows: (1000 × A_420_) ÷ (t × v × OD_600_), where t is min and v is ml.

### Gradient plate assay

Gradient plates were prepared as previously described (Bryson & Szybalski, 1952) with slight modifications. In brief, a 100 mm x 100 mm x 1.5 mm square petri dish (Thomas Scientific) was propped up on a 1 cm ledge to create a slant. 40 ml of LB with 1.5% agar and either 10 µg/ml chloramphenicol or 300 µg/ml erythromycin was poured and allowed to harden. The plate was then laid flat and 40 ml of LB with 1.5% agar was overlaid. For strains carrying plasmids, both top and bottom agar were also supplemented with 50 µg/ml carbenicillin and 0.02% arabinose. Strains were grown in LB (with carbenicillin for plasmid-containing strains) at 37°C with shaking at 250 rpm to exponential phase (OD_600_∼0.4), and 15 µl was dripped down the plate starting from the edge with the lowest concentration. The drips were allowed to dry and the plates were incubated at 37°C for 16 h and imaged using the fluorescein setting on a ChemiDoc MP Imaging machine (Biorad).

## Acknowledgements

The work was supported by an ERC advanced award (742210) and by a Wellcome Trust Investigator award to BFL. CN was supported by a Gates-Cambridge Scholarship and AN by a Herchel Smith Research Studentship. Work by LMR, MWO and GS was supported by the Intramural Program of the Eunice Kennedy Shriver National Institute of Child Health and Human Development. We acknowledge Diamond for access and support of the Cryo-EM facilities at the UK national electron bio-imaging centre (eBIC), funded by the Wellcome Trust, MRC and BBSRC. We thank colleagues at eBIC, especially Dan Clare and Alistair Siebert, for help with cryo-EM data collection. We thank Dima Chirgadze for help with data collection using the University of Cambridge cryo-EM facility. We also thank Douglas B. Weibel for kindly providing strains deficient in cardiolipin biogenesis. We are extremely grateful to Tristan Croll for advice and help with ISOLDE for the refinement of the models, and to Anirban Banerjee and Alex Sodt for helpful and insightful discussions.

## Author contributions

Studies were planned by D.D., A.N., M.W.O., C.E.N, P.-C.H., F.S., A.S.-H., L.M.R., M.D., S.K., G.S. and B.F.L. and carried out by D.D., A.N., M.W.O., C.E.N, P.-C.H., F.S., A.S.-H., L.M.R., M.D. and S.K. D.D., A.N., M.W.O., C.E.N, P.-C.H., F.S., A.S.-H., L.M.R., M.D., S.K., G.S. and B.F.L. carried out data analysis and manuscript preparation.

## Supporting Information

Appendix

